# Multi-channel intraneural vagus nerve recordings with a novel high-density carbon fiber microelectrode array

**DOI:** 10.1101/2020.05.15.098301

**Authors:** Ahmad A. Jiman, David C. Ratze, Elissa J. Welle, Paras R. Patel, Julianna M. Richie, Elizabeth C. Bottorff, John P. Seymour, Cynthia A. Chestek, Tim M. Bruns

**Affiliations:** Department of Biomedical Engineering, University of Michigan, Ann Arbor, MI, USA; Biointerfaces Institute, University of Michigan, Ann Arbor, MI, USA; Department of Electrical and Computer Engineering, King Abdulaziz University, Jeddah, Saudi Arabia; Department of Electrical Engineering and Computer Science, University of Michigan, Ann Arbor, MI, USA; Department of Neurosurgery, University of Texas Health Science Center, Houston, TX, USA

## Abstract

Autonomic nerves convey essential neural signals that regulate vital body functions. Recording clearly distinctive physiological neural signals from autonomic nerves will help develop new treatments for restoring regulatory functions. However, this is very challenging due to the small nature of autonomic nerves and the low-amplitude signals from their small axons. We developed a multi-channel, high-density, intraneural carbon fiber microelectrode array (CFMA) with ultra-small electrodes (8-9 μm in diameter, 150-250 μm in length) for recording physiological action potentials from small autonomic nerves. In this study, we inserted CFMA with up to 16 recording carbon fibers in the cervical vagus nerve of 22 isoflurane-anesthetized rats. We recorded action potentials with peak-to-peak amplitudes of 15.1-91.7 μV and signal-to-noise ratios of 2.0-8.3 on multiple carbon fibers per experiment, determined conduction velocities of some vagal signals in the afferent (0.7-4.4 m/sec) and efferent (0.7-8.8 m/sec) directions, and monitored firing rate changes in breathing and blood glucose modulated conditions. Overall, these experiments demonstrated that CFMA is a novel interface for in-vivo intraneural action potential recordings. This work is considerable progress towards the comprehensive understanding of physiological neural signaling in vital regulatory functions controlled by autonomic nerves.

## Introduction

The autonomic nervous system has a major role in the regulation of unconscious functions that are essential to the body. The system is divided into the sympathetic nervous system, which controls “fight-or-flight” responses, and the parasympathetic nervous system, which regulates “rest-and-digest” functions^1^. A main parasympathetic nerve is the vagus nerve, which innervates many visceral organs, such as the heart, lungs, stomach, liver, pancreas and intestines^2,3^, and contributes to the regulation of numerous autonomic functions, which include breathing, immune responses, digestion, glucose metabolism and others^4–8^. The vagus nerve at the cervical level is partially composed of myelinated Aδ and B fibers^9,10^, but the great majority of axons (over 80%) are unmyelinated C-fibers^2,11,12^. These fibers predominantly convey afferent (sensory) signals from the innervated organs to the central nervous system^13^. Hence, the vagus nerve is an attractive target for monitoring the physiological state of visceral organs for therapeutic or scientific objectives.

A class of therapies that has gained considerable interest in recent years is bioelectronic medicine, which targets autonomic nerves to detect and alter neural activity for restoring autonomic functions^14–16^. The variety of bioelectronic medicine applications that target the vagus nerve have led to clinical trials on vagus nerve stimulation (VNS) for patients with epilepsy^17^, stroke^18^, depression^19^, rheumatoid arthritis^20^, obesity^21^, and type-2 diabetes^22^, among others. Despite the therapeutic benefits of VNS and bioelectronic medicine, stimulation patterns are generally selected by experimenting with different parameters without monitoring the physiological signaling in the nerve. A key element that is needed to achieve the full potential of bioelectronic medicine is a better understanding of neural signaling in normal and modulated physiological conditions.

Recording neural activity from autonomic nerves is very challenging due to the often sub-millimeter nature of these nerves^23^, the protective layers surrounding the nerve (epineurium), bundle of axons (perineurium) and individual axons (endoneurium)^24,25^, and the low-amplitude waveforms generated from small unmyelinated C-fibers^26^ that dominate autonomic nerves^27,28^. Studies have applied electrical stimulation on autonomic nerves to record evoked neural activity using extraneural electrodes, which record from outside the nerve^9,29^, and intraneural electrodes, which penetrate the nerve^30^. Although electrical stimulation-evoked responses can be useful in determining the type of activated fibers, these responses do not represent physiological neural signaling. A few research groups have obtained physiological neural recordings from autonomic nerves using extraneural cuff electrodes^31–35^. However, extraneural electrodes lack spatial selectivity, as these electrodes record the compound activity of hundreds to thousands of axons from outside the nerve. Intraneural electrodes penetrate the nerve to be closer to axons and provide better selectivity and higher signal-to-noise ratio (SNR) recordings than extraneural electrodes^24,25^. Intraneural high-density Utah slanted electrode arrays (HD-USEAs) have 48 electrodes (30-100 μm tapered diameter) in a 5×10 configuration (pitch of 200 μm; corner electrodes used as reference and ground) and have been used to record signals in cat pudendal nerves, which have an approximate diameter of 1 mm^36,37^. This configuration of the silicon-based HD-USEA is large (1×2 mm) and rigid for most autonomic nerves, which are often under 1 mm in diameter. For recording from small-diameter (≤ 0.5 mm) autonomic nerves, carbon nanotube (CNT) electrodes have demonstrated high SNR recordings (> 10 dB) in rat glossopharyngeal and vagus nerves (diameter of 100-300 μm)^23^. This was achieved by inserting two CNT electrodes (10 μm in diameter) in a nerve target at a 2-mm separation to obtain a single differential recording. Another research group inserted 4-channel carbon fiber arrays (electrode diameter ≤ 15 μm, pitch of 150 μm) in tracheosyringeal nerves of zebra finch birds, which are 125 μm in diameter and mostly composed of myelinated fibers (99%)^30^. They obtained spontaneous recordings but primarily demonstrated electrical stimulation-evoked compound neural responses. Autonomic nerves are typically dominated by hundreds to thousands of unmyelinated fibers^27,28,38,39^. The fibers of the vagus nerve in particular innervate multiple critical organs and contribute to the regulation of many autonomic functions^2–8^. Therefore, a need remains for an intraneural electrode array that can record physiological single-neuron activity at multiple sampling locations within small autonomic nerves.

Our research group has developed a novel, multi-channel, blowtorch-sharpened, intraneural carbon fiber microelectrode array (CFMA) for small autonomic nerves. The CFMA has ultra-small recording electrodes (8-9 μm in diameter, 150-250 μm in length) in a higher-density configuration than previously reported arrays (16 carbon fibers in 2 rows, with a 132 μm pitch along the array and 50 μm between rows). Prior versions of the CFMA with longer (500-5000 μm), unsharpened carbon fibers have demonstrated high SNR recordings with minimal tissue damage in the rat cerebral cortex^40–43^. We hypothesized that this novel CFMA would obtain physiological action potential recordings with high SNR in small autonomic nerves. In this study, we inserted CFMAs in rat cervical vagus nerves (diameter of 300-500 μm). We recorded action potentials on multiple carbon fibers per experiment, determined the propagation direction and conduction velocity of some vagal signals, and monitored changes in neural activity in breathing and blood glucose modulated conditions.

## Results

We fabricated CFMAs with 16 carbon fibers in a 2×8 configuration. Fibers within each row had a pitch of 132 μm, and the two rows were separated by 50 μm (Fig. 1a). The carbon fibers, which had a diameter of 8-9 μm, were cut to 150-250 μm in length and the tips were sharpened with a blowtorch (Fig. 1b). The active recording site for a carbon fiber is coated with poly(3,4-ethylene-dioxythiophene):sodium p-toluenesulfonate (PEDOT:pTS) and spans 135-160 μm in length from the tip. To facilitate CFMA insertion in a rat vagus nerve, we designed a nerve-holder to secure and elevate the vagus nerve away from fluid and breathing motions of the cervical cavity, and allow accurate positioning of a small camera to visualize the CFMA-nerve interface during insertion (Fig. 1c).

**Figure 1.**
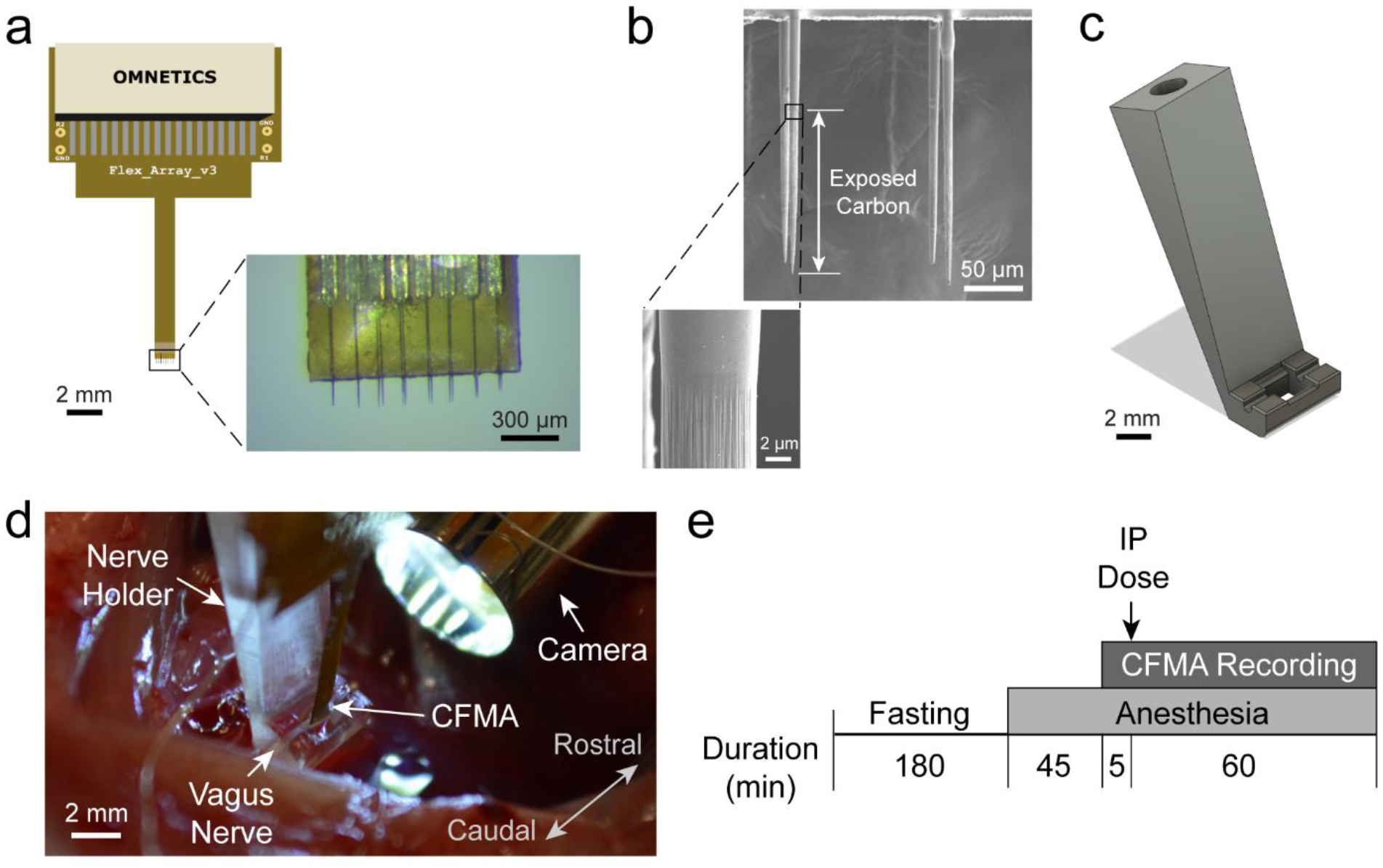
Carbon Fiber Microelectrode Array (CFMA) and experimental setup. **(a)** CFMA with 16 blowtorch-sharpened carbon fibers in a 2×8 configuration. **(b)** Scanning Electron Microscopy (SEM) images of sharpened carbon fibers. Arrows indicate an exposed (non-insulated) carbon fiber region with a length of ~140 μm from the tip. **(c)** Design of a nerve-holder to facilitate CFMA insertion in a rat vagus nerve. Dimensions of the design are shown in Supplementary Fig. S3. **(d)** Surgical setup for inserting CFMA in the vagus nerve. **(e)** Timeline for the experimental protocol. An intraperitoneal (IP) dose of glucose, insulin, 2-deoxy-D-glucose, or saline was injected to modulate blood glucose concentration levels.

We inserted 6 CFMAs in the left cervical vagus nerve of 22 Sprague-Dawley rats. We observed neural activity on 167 out of 326 inserted functional carbon fibers (impedance < 1 MΩ). The neural activity on each carbon fiber was sorted into 1 neural cluster (n=160) or 2 neural clusters (n=7). The functional carbon fibers had an average impedance of 31.3 ± 42.0 kΩ (mean ± standard deviation) in saline before an experiment, 70.8 ± 81.9 kΩ in the nerve immediately after insertion, and 94.7 ± 146.7 kΩ in the nerve at the end of the experiment. The variability in impedance is likely due to the tip sharpening process, which leads to varying active recording site lengths from the tip. Three of the CFMAs were used in more than one experiment (4-8 experiments per CFMA), which initially had a total of 48 functional carbon fibers (16 carbon fibers per CFMA) with an average impedance of 52.8 ± 36.8 kΩ after insertion in the first experiment. After insertion in the fourth experiment, 45 carbon fibers on these three CFMAs (14-16 carbon fibers per CFMA) remained functional with an average impedance of 92.6 ± 149.5 kΩ. On average for a single experiment, we made 2.3 ± 2.9 attempts to insert a CFMA with 14.8 ± 1.8 functional carbon fibers and observed neural activity on 7.6 ± 5.8 carbon fibers. In 59% of the experiments, only one insertion attempt was needed to observe neural activity. If no neural activity was observed, another 1-3 insertion attempts were made at the same location before moving to another location on the nerve. There were no distinctive differences among the recordings of rats with different gender or sizes.

### Multi-Channel Recordings of Vagal Nerve Activity

We observed physiological neural activity in the vagus nerve on at least one recording carbon fiber in 19 of the total 22 experiments. The recorded neural activity was sorted into clusters and the mean peak-to-peak amplitudes of the sorted clusters were between 15.1 and 91.7 μV with SNR of 2.0-8.3. An example of vagal nerve activity on multiple recording carbon fibers from the same experiment is shown in Fig. 2a. Another example showing neural activity on all 16 recording carbon fibers is shown by Supplementary Fig. S1.

**Figure 2.**
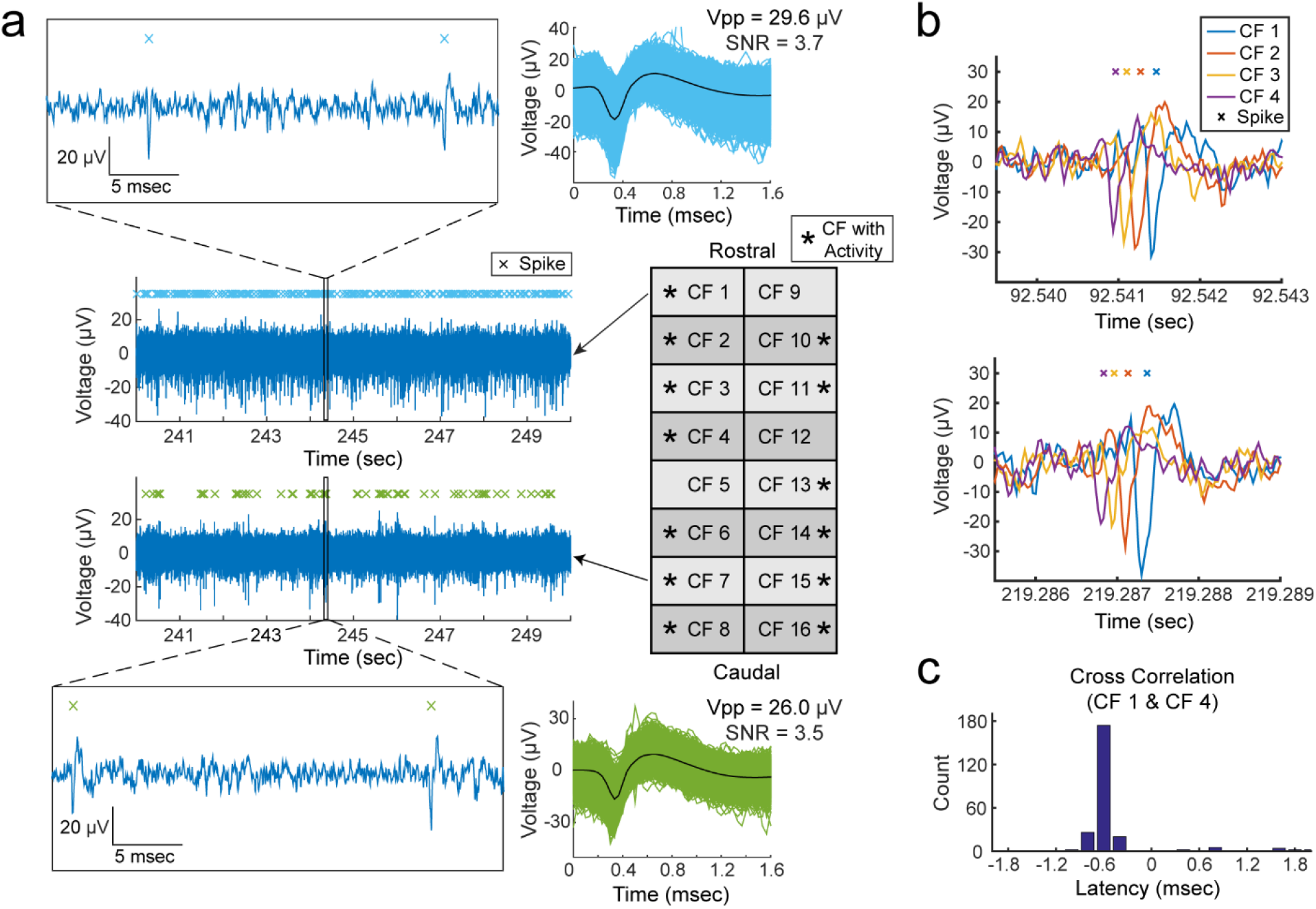
Representative recordings of physiological vagal nerve activity and signal propagation along CFMA carbon fibers (CFs). **(a)** Recordings of vagal nerve activity on 2 carbon fibers (CF 1 and CF 7) in the same experiment showing distinctive signals. The firing of sorted spikes (marked with x) are unique across CFs. **(b)** Instances of signal propagation along CF 1 – CF 4. **(c)** Cross correlation of spikes on CF 1 and CF 4. The prevalent latency occurred at −0.6 msec with a count of 174 spikes (55.8% of spikes on CF 4). At latency of −0.6 msec, the spikes occurred on CF 4 before CF 1, suggesting that the signal is propagating in the afferent direction at a conduction velocity of 0.7 m/sec.

Propagation of vagal signals were detected along adjacent recording carbon fibers in some experiments. We observed neural signals in 10 experiments propagating in the afferent direction with conduction velocities of 0.7-4.4 m/sec over the span of 2-7 carbon fibers (132-792 μm). Furthermore, we monitored efferent signals conducting at 0.7-8.8 m/sec along 2-5 carbon fibers (132-528 μm) in 5 experiments. Examples of propagating afferent signals are shown in Fig. 2b with cross-correlation to inspect the latency of those signals along CFMA carbon fibers (Fig. 2c). Neural activity of carbon fibers on opposite sides of each other (50 μm apart) were generally independent. In some instances (n=37), opposite carbon fibers recorded coinciding spikes with no time delay but with different amplitudes, suggesting that these spikes are generated from a neuron located between these opposite carbon fibers. Although afferent signals in 7 experiments and efferent signals in 3 experiments were detected propagating along adjacent carbon fibers on both rows of a CFMA, these propagating signals on opposite rows were independent of each other. A representative experiment that demonstrated these signals is shown in Supplementary Fig. S2.

### Breathing-Related Neural Activity

We observed vagal signals with periodic bursting firing behavior (n=6 experiments) at repetition rates of 39.4 ± 10.8 cycles/min, which were similar to the animals’ breathing rates of 39.3 ± 9.9 breaths/min (Fig. 3). In a subset of experiments (n=3), we reduced the breathing rate to 20.0 ± 8.0 breaths/min by increasing the depth of anesthesia, and the firing-burst repetition rates reduced to a similar level at 19.1 ± 10.3 cycles/min, with maintained peak-to-peak amplitudes (31.7 ± 11.6 μV to 29.3 ± 10.1 μV) and inter-spike interval (ISI) peak values (9.2 ± 1.6 msec to 10.5 ± 1.6 msec), as shown in the example in Fig. 3a. The periodic bursting behaviors were usually firing at one ISI peak of 9.5 ± 1.3 msec. However, in two experiments, two distinct ISI peaks were observed at 9.8 ± 1.8 msec and 24.2 ± 6.0 msec (e.g. Fig. 3b). The occurrence of these breathing-related signals on only 1-2 channels out of 14-16 functional channels (impedance < 1 MΩ) per experiment strongly suggests that these are neural signals related to breathing, rather than motion artifacts which would have occurred on all functional channels inserted in the same region of a nerve (i.e. cervical vagus nerve) as they encounter the same breathing motion.

**Figure 3.**
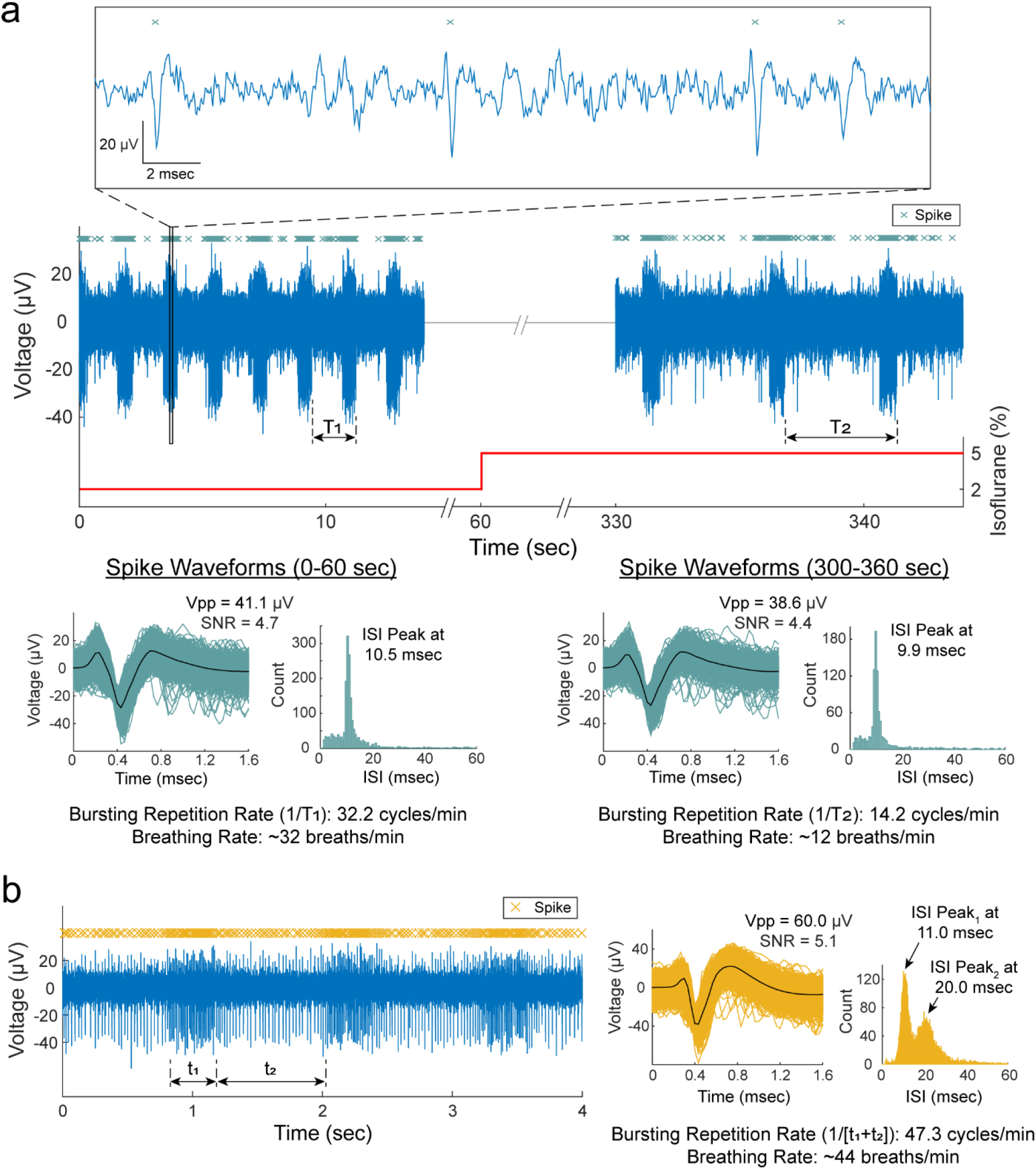
Breathing-related neural activity. **(a)** Recordings of vagus nerve activity at 2% and 5% isoflurane. The bursting firing behavior had a repetition rate of 32.2 cycles/min (1/T1) during animal’s breathing rate of ~32 breaths/min at 2% isoflurane. The firing-behavior repetition rate reduced to 14.2 cycles/min (1/T2) as the breathing rate reduced to ~12 breaths/min at 5% isoflurane. The average peak-to-peak amplitude (Vpp) and inter-spike interval (ISI) peaks were similar for the spike waveforms at 2% and 5% isoflurane. **(b)** Bursting firing behavior at two distinct ISIs indicated at the t1 and t2 durations. The animal’s breathing rate was ~44 breaths/min and the repetition rate for the bursting firing behavior was 47.3 cycles/min (1/[t1+t2]).

### Neural Firing Rate Behavior in Blood Glucose Modulation Conditions

In each experiment, we recorded vagal nerve activity for a baseline period of at least 5 minutes before the intraperitoneal (IP) injection of a blood glucose modulation dose (glucose, insulin, 2-deoxy-D-glucose, or saline). Recordings were continued for 60 minutes after the injection. Physiological parameters (blood glucose concentration, breathing rate, and heart rate) were measured every 5 minutes throughout the entire experiment. The recorded neural activity were sorted into 174 clusters. The firing rate of these clusters showed moderate or high correlation coefficients (|ρ| ≥ 0.3) with one (n=15), two (n=35), or all three (n=96) of the tracked physiological parameters. However, correlation coefficients did not show clear associations between any glucose modulation dosing and any of the physiological parameters across experiments. An experiment with a carbon fiber that recorded the activity of 2 sorted clusters, along with the physiological measurements (blood glucose concentration, breathing rate and heart rate) and correlation coefficients, is shown in Fig. 4.

**Figure 4.**
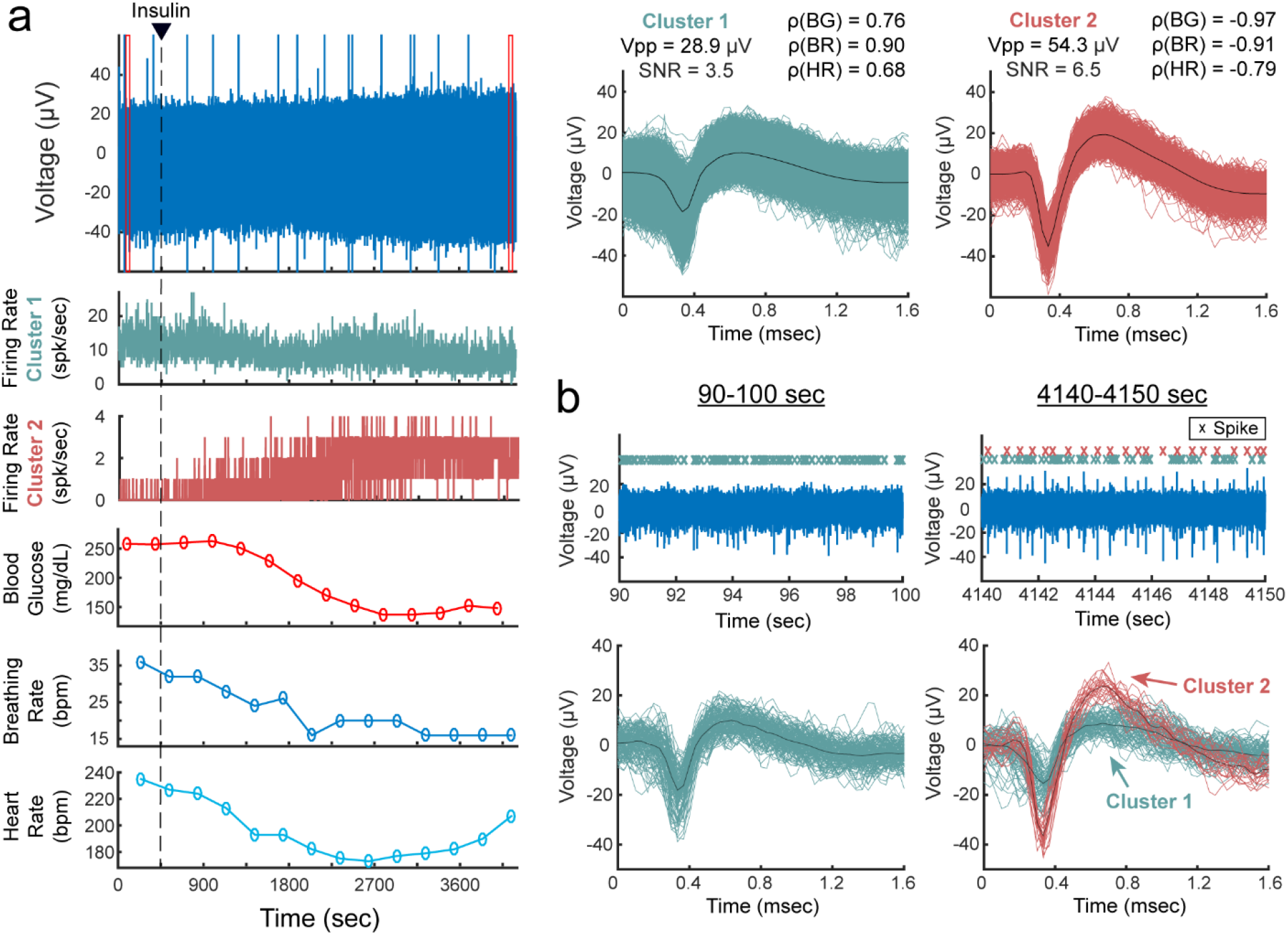
Vagus nerve recordings with sorted clusters in an insulin-injected experiment. **(a)** Filtered signal of a vagus nerve recording, firing rates of the two sorted clusters, and measurements of blood glucose (BG) concentration’ breathing rate (BR) and heart rate (HR). The waveforms of the sorted clusters are shown with the average peak-to-peak amplitude (Vpp), signal-to-noise ratio (SNR), and correlation coefficients (ρ) between cluster firing rate and BG, BR and HR measurements. The red boxes in the voltage plot indicate the time-window for the plots in b. The large amplitude spikes in the voltage plot are noise artifacts, which we confirmed by visually identifying the non-action potential shape of their waveforms (not shown in figure). **(b)** Filtered signal of the vagus nerve recording before and after insulin injection, and the spike waveforms of the two sorted clusters within each 10-sec window.

Although correlation coefficients did not show a clear relationship between any of the glucose modulation doses and physiological parameters, we observed clusters with interesting firing rate behaviors after injection of a modulation dose, as shown in Fig. 5. In some glucose injection experiments (n=4, 66.7%), we observed neural clusters (n=11, 36.7% from all glucose experiments) with an average peak-to-peak amplitude of 24.7 ± 6.4 μV and initial firing rate of 6.8 ± 8.9 spikes/sec that decreased after administration of glucose to 1.8 ± 2.4 spikes/sec. The clusters in one of these experiments showed signal propagation in the afferent direction at a conduction velocity of 0.7 m/sec. In some experiments with an insulin injection (n=4, 66.7%), neural clusters (n=4, 6.8% from all insulin experiments) with amplitudes of 53.3 ± 28.0 μV peak-to-peak increased their firing rates from 1.2 ± 1.8 spikes/sec to 6.0 ± 8.3 spikes/sec at 1-13 minutes after insulin administration. Injection of 2-deoxy-D-glucose (2-DG) induced a similar neural response to insulin in some experiments (n=2, 33.3%). Starting at 1-9 minutes after 2-DG administration, clusters (n=6, 10.7% from all 2-DG experiments) with an average amplitude of 29.2 ± 8.1 μV peak-to-peak increased their firing rates from 3.8 ± 4.6 spikes/sec to 7.6 ± 8.2 spikes/sec. Two of these clusters that responded to 2-DG were propagating in the afferent direction at a conduction velocity of 0.7 m/sec. The clusters from all the experiments with a saline injection (n=29) had an average peak-to-peak amplitude of 38.4 ± 11.9 μV. The firing rate was 4.3 ± 9.8 spikes/sec before the saline injection and 2.7 ± 5.6 spikes/sec after the injection. A summary of all the performed experiments is shown in Table 1.

**Figure 5.**
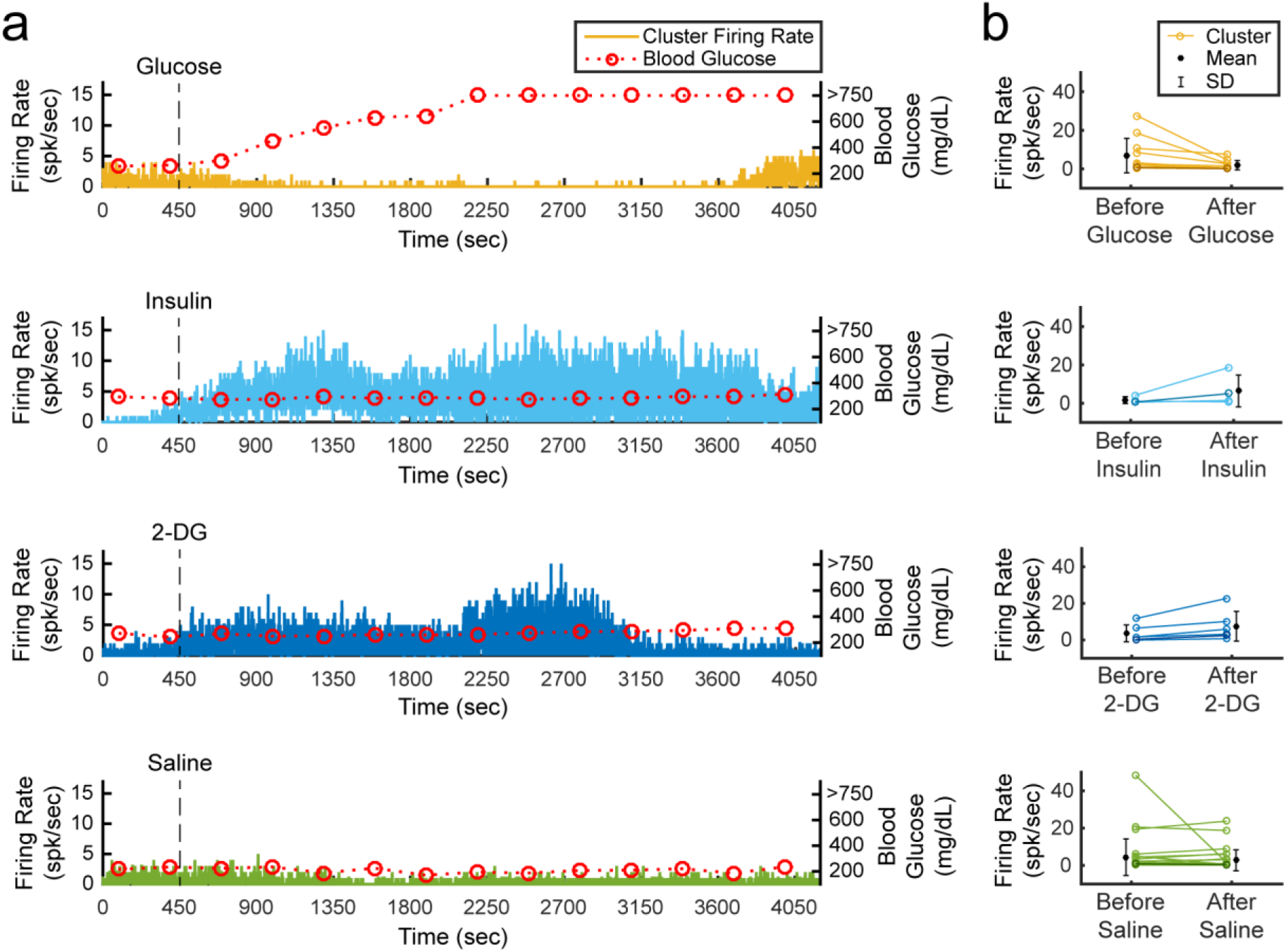
Cluster firing rates in blood glucose modulated conditions. **(a)** Firing rate behavior of representative clusters observed in experiments with an intraperitoneal (IP) injection of glucose, insulin, 2-deoxy-D-glucose (2-DG), or saline. Blood glucose concentration measurements above 750 mg/dL were not available due to the limitations of the glucometer. **(b)** Changes in firing rates for all the clusters that responded to glucose, insulin and 2-DG injections, and all the clusters observed in experiments with a saline injection. An injection was administered at 450 sec and the mean cluster firing rate was calculated before the injection (0-400 sec) and after the injection (500-3000 sec). Clusters with darkened colors are the representative clusters in panel a.

**Table 1.** Summary of all the experiments with carbon fiber microelectrode array (CFMA) inserted in the vagus nerve.

| ID | Array | Inserted Functional CF | CF with Neural Activity | Sorted Clusters | Vpp Range ($\mu\text{V}$) | SNR Range | IP Dose | Responding Clusters | | Observed Signal Propagation Range (m/sec) | | |
| --- | --- | --- | --- | --- | --- | --- | --- | --- | --- | --- | --- | --- |
| | | | | | | | | IP Dose | Breathing | Afferent (n) | Efferent (n) | Max Span ($\mu\text{m}$) |
| 1 | A | 16 | 16 | 16 | 21.8 - 27.5 | 2.4 - 3.1 | Glucose | 6 | 0 | - | - | - |
| 2 | A | 16 | 6 | 6 | 18.7 - 34.5 | 2.3 - 3.5 | 2-DG | 0 | 0 | - | - | - |
| 3 | A | 15 | 5 | 5 | 17.9 - 29.2 | 2.0 - 3.0 | Glucose | 1 | 0 | - | - | - |
| 4 | A | 15 | 14 | 15 | 23.2 - 91.7 | 2.5 - 8.3 | Insulin | 1 | 1 | 0.7 (2) | 0.7 (2) | 396 |
| 5 | A | 15 | 7 | 8 | 22.0 - 43.9 | 2.5 - 3.5 | Saline | - | 0 | 0.7 (2) | 0.7 (2) | 264 |
| 6 | A | 14 | 0 | 0 | - | - | Insulin | 0 | 0 | - | - | - |
| 7 | B | 16 | 0 | 0 | - | - | Glucose | 0 | 0 | - | - | - |
| 8 | C | 16 | 7 | 8 | 20.4 - 40.5 | 2.9 - 5.3 | Insulin | 1 | 1 | - | - | - |
| 9 | C | 16 | 6 | 7 | 20.7 - 59.1 | 3.3 - 7.2 | Saline | - | 1 | - | 0.7 (1) | 132 |
| 10 | C | 16 | 16 | 16 | 21.1 - 58.7 | 3.0 - 6.9 | 2-DG | 0 | 1 | 0.7-4.4 (3) | 0.7 - 8.8 (6) | 528 |
| 11 | D | 16 | 6 | 6 | 15.1 - 38.4 | 2.8 - 4.6 | Glucose | 3 | 0 | 0.7 (1) | - | 132 |
| 12 | E | 14 | 5 | 6 | 17.7 - 78.2 | 2.8 - 7.9 | Insulin | 1 | 1 | - | - | - |
| 13 | C | 14 | 2 | 2 | 27.5 - 44.0 | 3.6 - 5.7 | 2-DG | 0 | 0 | - | - | - |
| 14 | A | 14 | 14 | 14 | 27.4 - 62.8 | 2.7 - 5.5 | Saline | - | 0 | 0.7-0.9 (3) | - | 792 |
| 15 | D | 16 | 14 | 16 | 18.7 - 54.3 | 2.4 - 6.5 | Insulin | 1 | 0 | 0.7 (3) | - | 660 |
| 16 | D | 16 | 13 | 13 | 17.3 - 35.6 | 2.4 - 4.0 | 2-DG | 4 | 2 | 0.7 (3) | - | 792 |
| 17 | D | 16 | 14 | 14 | 20.0 - 26.6 | 2.8 - 3.5 | Insulin | 0 | 0 | 0.7-1.0 (5) | - | 528 |
| 18 | D | 15 | 2 | 2 | 18.8 - 26.0 | 3.1 - 3.2 | Glucose | 1 | 0 | 0.7 (1) | - | 132 |
| 19 | D | 15 | 4 | 4 | 19.3 - 36.5 | 3.1 - 4.7 | 2-DG | 0 | 0 | - | - | - |
| 20 | F | 8 | 0 | 0 | - | - | Saline | - | 0 | - | - | - |
| 21 | A | 12 | 1 | 1 | 26.7 | 2.8 | Glucose | 0 | 0 | - | - | - |
| 22 | D | 15 | 15 | 15 | 27.5 - 54.0 | 3.1 - 6.4 | 2-DG | 2 | 0 | 0.7 - 0.9 (6) | 0.7 (1) | 528 |
| Total |  | 326 | 167 | 174 | 15.1 - 91.7 | 2.0 - 8.3 | - | 21 | 7 | 0.7 - 4.4 | 0.7 - 8.8 | 132 - 792 |

## Discussion

We developed a multi-channel, high-density, intraneural carbon fiber microelectrode array (CFMA) for recording neural signals in autonomic nerves (Fig. 1). Using the CFMA, we obtained axonal action potential recordings in rat cervical vagus nerves with signal-to-noise ratio (SNR) of 2.0-8.3. We recorded physiological vagal nerve activity that was unique across multiple channels per experiment (Fig. 2a), determined the propagation direction and conduction velocity of some vagal signals (Fig. 2b), and monitored changes in neural activity in physiologically modulated conditions (Figs. 3-5). These data demonstrate CFMA as a new interface for in-vivo intraneural recordings. This work, to our knowledge, is the first to demonstrate in-vivo physiological action potential recordings on multiple channels in a sub-millimeter autonomic nerve. Monitoring physiological signaling in autonomic nerves will help researchers better understand the neural control and feedback processes for autonomic organs, which is a key element for developing innovative treatment modalities to restore vital body functions regulated by autonomic nerves.

Our experimental recordings demonstrated CFMA as a multi-channel, intraneural array for small-diameter (≤ 0.5 mm) autonomic nerves. In prior work, the high-density Utah slanted electrode array (HD-USEA) was implanted in a 1-mm diameter cat pudendal nerve^36^. While the 48-channel, 200 μm pitch HD-USEA (footprint over 1 mm^2^) was used to record physiological signaling from autonomic organs (feline lower urinary tract)^36^, the size of the HD-USEA electrode shanks (300-800 μm in length, 30-100 μm in diameter)^37^ are much larger than in the CFMA and would make intraneural recordings in small autonomic nerves challenging. In our study, 16-channel CFMAs (footprint less than 0.05 mm^2^; 132 μm pitch along the array and 50 μm between two rows) with ultra-small electrodes (150-250 μm in length, 8-9 μm in diameter) were implanted in small-diameter (300-500 μm) rat vagus nerves. Another intraneural electrode that obtained physiological recordings in small-diameter (100-300 μm) autonomic nerves (rat glossopharyngeal and vagus nerves) are carbon nanotube (CNT) electrodes^23^. Two single-channel CNT electrodes were inserted with a 2-mm separation in a nerve target to obtain only a single differential recording in that study. The CFMA recorded physiological neural activity on multiple channels (up to 16 channels), which allowed us to detect the propagation direction and conduction velocity of some signals (Fig. 2). The recording exposure site on each carbon fiber spans 135-160 μm in length from the tip, which provided better spatial selectivity recordings than CNT electrodes that had an exposed recording segment of ~500 μm. Another research group developed an intraneural 4-channel carbon fiber array with a similar pitch (150 μm) as the CFMA but with longer carbon fibers (≥ 350 μm) that recorded from tracheosyringeal nerves of zebra finch birds (diameter of 125 μm)^30^. They demonstrated an innovative blowtorching technique for sharpening carbon fibers to directly insert carbon fibers in a nerve, which we adapted to our shorter (150-250 μm) CFMA carbon fibers. Although an example of spontaneous activity was shown using the 4-channel array, the majority of their demonstrated signals were evoked responses from electrical stimulation.

The observed spike waveforms in CFMA recordings from the vagus nerve (e.g. Figs. 2-5) are action potentials generated by individual neurons, based on the waveform shape and time scale (1-2 msec)^44,45^. Furthermore, we observed propagation of signals in the afferent and efferent direction within the conduction velocity range for myelinated (Aδ and B) and unmyelinated (C) fibers (e.g. Fig. 2 and Supplementary Fig. S2), which are present in the vagus nerve^9,10^. However, due to the similarity in the waveform shapes and the normal variations in waveform amplitudes, we were not able to sort the detected action potentials into clear single units per channel, with only a few channels yielding more than one sortable cluster. The active recording site for a CFMA carbon fiber spans 135-160 μm in length from the tip (Fig. 1), which exposes the recording site to an estimation of over 200 axons within a distance of 5 μm from the recording site (Fig. 6). This estimation is based on the approximate axon density in the rat vagus nerve, which has around 11,000 axons^27,28^ contained within a diameter of about 300 μm. Further work on reducing the exposed recording site area may assist in monitoring more localized axon activity with a lower background noise level^41,43,46^. Moreover, current spike-sorting algorithms are mostly designed for central nervous system recordings^47^, which assume the waveforms are from neuron cell bodies that generate higher amplitude waveforms and have more diverse shapes than unmyelinated axons. Future work is needed to study the recording nature in autonomic nerves and develop spike-sorting algorithms for axonal recordings.

**Figure 6.**
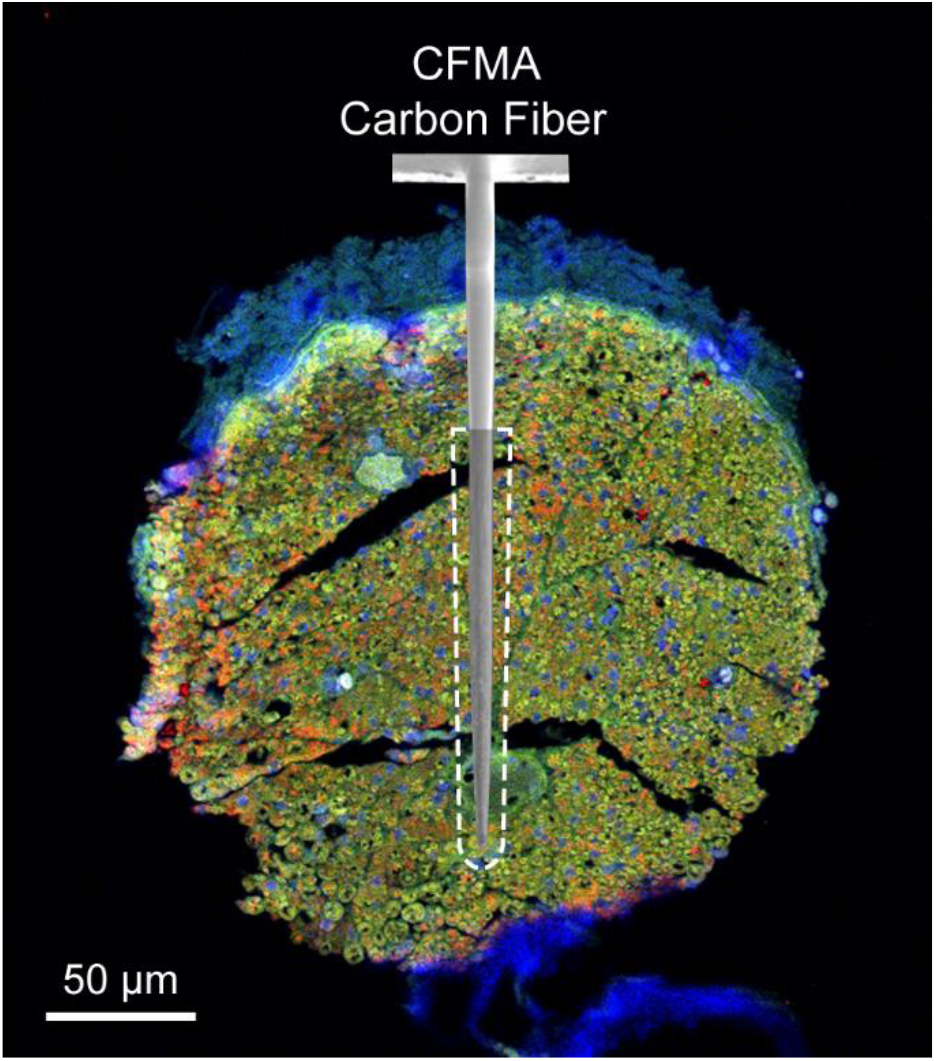
Diagram of a CFMA carbon fiber overlaid on an immunohistochemistry image of a rat cervical vagus nerve. The shaded region on the CFMA carbon fiber is the active recording site. The dashed lines specify an area within a distance of 5 μm from the active recording site of one carbon fiber. The area is estimated to be occupied by over 200 axons based on the typical axon density of a rat vagus nerve (~11,000 axons in 300 μm diameter nerve)^27,28^. The nerve sample in this image was stained with 4’, 6-diamino-2-phenylindole (DAPI), myelin basic protein (MBP), and anti-beta III tubuline (TUJ1) to show nucleotides (blue), myelin (green) and axons (red), respectively.

Sorted clusters from our recorded vagal nerve activity showed interesting firing rate behavior that may be related to the measured physiological parameters of breathing rate, heart rate and blood glucose concentrations. We observed neural clusters with periodic firing-burst behavior at repetition rates similar to the measured breathing rates (Fig. 3). Vagus nerve fibers innervate the lungs, with critical relevance for breathing control^4,48,49^. Our observed vagal signals may be related to the neural control over breathing or an afferent response to chest and/or lung expansion^48^. We also observed interesting changes in vagal firing rate behavior after injection of blood glucose modulation doses (Fig. 5). These experiments were performed on fasted rats, and neural signals before the dose injection may represent vagal afferent signals to drive an increase in glucose intake. The firing rate of clusters decreased after administration of glucose, which may suggest that the signaling for glucose intake was met. This observation aligns with previous studies that showed neural recordings from dissected fibers of afferent hepatic vagus nerve branches in isolated and perfused livers using wire electrodes^50^, and compound neural activity from the cervical vagus nerve using cuff electrodes after removing the nerve sheath^35^. Both studies demonstrated similar firing rate changes following the administration of glucose. The CFMA was also able to determine the propagation of some of these signals in the afferent direction at a conduction velocity for unmyelinated C-fibers, which dominate the vagus nerve^27,28^. The increase in firing rate observed after insulin or 2-deoxy-D-glucose (2-DG) injection, which induce similar symptoms, may represent a surge of afferent activity to enhance the request for glucose intake, which aligns with the previously mentioned studies that also showed increased afferent activity in the hepatic branch of vagus nerves using wire electrodes^50^, and increased compound activity of cervical vagus nerves using cuff electrodes^35^ within 10 minutes after administration of insulin or 2-DG. Similarly, the CFMA detected the afferent propagation of some of these signals at a C-fiber conduction velocity. However, these observed responses were inconsistent across our experiments with identical injection doses. Only 66.7% of the experiments yielded clusters that responded to a glucose or insulin dose, compared to a previous study that detected responding clusters in 83.3% of the performed glucose and insulin experiments using cuff electrodes placed on the cervical vagus nerve of mice^35^. This may be due to variations in CMFA sampling of neural activity within the vagus nerve across experiments while cuff electrodes record the overall nerve activity. However, CFMA records action potentials generated from individual neurons whereas a cuff electrode records the compound neural activity of the entire nerve, potentially allowing for final signal resolution. Moreover, there were similarities in the blood glucose concentration trends during an experiment to the other measured physiological parameters (i.e. breathing rate and heart rate) in most experiments (e.g. Fig. 4). The anesthetic agent we used in our experiments was isoflurane, which maintained consistent and stable depth of anesthesia for recording vagal nerve activity with ultra-small carbon fibers. In preliminary experiments using other agents (e.g. ketamine), occasional muscle twitches would lead to CFMA movement or carbon fiber breakage due to the CFMA being attached to an externally-mounted inserter. These muscle twitches were not observed under isoflurane anesthesia. However, isoflurane suppresses neural activity in the central and autonomic nervous systems and impacts multiple physiological parameters, including blood glucose concentration, respiration, and arterial pressure^51,52^. This may explain the substantial changes observed in breathing rate and heart rate (e.g. Fig. 4) that were likely incidental changes caused by isoflurane anesthesia maintained at a constant rate for long durations (more than 1 hour)^51,52^ which may have led to the moderate or high correlation coefficients between cluster firing rates and the measured physiological parameters. As we mentioned earlier, the identified clusters are the combined recordings of action potentials from multiple neurons conveying different information (Fig. 6). While the activity of many of these individual axons may not be correlated with the measured physiological parameters, the presence of at least a few axons related to the measured parameters in each cluster likely contributed to the consistent moderate or high correlations across all clusters. Although this work showed unique in-vivo action potential recordings from the vagus nerve, using more selective CFMA in experiments with minimal or no anesthesia would allow more physiological activities to occur and may be necessary to clearly link vagal nerve activity to physiological changes.

This work had numerous limitations. The exact insertion location for the CFMA arrays in the vagus nerve varied between our experiments. The rat cervical vagus nerve is estimated to contain around 11,000 axons^27,28^ that regulate many autonomic functions^4^. To illustrate this variation, potential breathing-related signals (e.g. Fig. 3) were only observed in 6 out of the 22 experiments, although all the rats were breathing normally during the experiments. Furthermore, we detected propagation direction of some, but not all, recorded vagal signals (e.g. Fig. 2), likely due to the variation in CFMA insertion alignment along the nerve. Additional work on redesigning the electrode configuration may be needed to cover a wider range of axonal activity while providing high selectivity for individual recording sites, such as with staggered rows of carbon fibers with variable lengths. Another limitation is the requirement to lift the nerve for CFMA insertion, which applies tension on the nerve due to the nerve-holder design. Although the nerve-holder added the risk of nerve injury, the nerve-holder was necessary to position a camera to visualize the alignment of CFMA carbon fibers with the vagus nerve for insertion (Fig. 1). Redesigning the nerve-holder and possibly restructuring the implantation procedure may be needed to eliminate the applied tension and avoid the risk of injuring the nerve. The potential nerve damage caused by CFMA insertion was not assessed in this study. Although prior versions of CFMA with longer, unsharpened carbon fibers showed minimal tissue damage in the rat cerebral cortex^40,42^, future studies with histological analysis are needed to assess the impact of CFMA in peripheral nerves. This study only demonstrated acute CFMA recordings from the rat vagus nerve. However, the CFMA has also recorded action potentials from the cat pudendal nerve and rat sural nerve, a branch of the sciatic nerve, in acute preliminary experiments (data not shown). The 50 μm pitch between the two rows of carbon fibers allows the CFMA to target smaller nerves, such as the mouse vagus nerve (~100 μm in diameter), which is another common animal model for vagus nerve interfacing studies^31,32,35^. While the CFMA design presented in this work is not suitable for chronic implants in nerves, sharpened carbon fibers could be combined with a soft substrate (e.g. silicone) to allow for strain relief in chronic implants^53^. Future acute and chronic studies on physiological neural recordings from various peripheral nerves of different animal models will provide new perspectives on neural control processes.

Overall, our experiments demonstrated that CFMA is a novel interface for in-vivo, high-density, multi-channel, intraneural action potential recordings in small autonomic nerves. Further work is needed to refine the selectivity of CFMA and develop a chronic form for long-term, behavioral recordings in autonomic nerves without the presence of anesthesia. This work provided insights in intraneural axonal recordings and is considerable progress towards the comprehensive understanding of physiological signaling in vital regulatory functions controlled by autonomic nerves.

## Methods

### Fabrication of Carbon Fiber Microelectrode Array

The independent components for fabricating carbon fiber microelectrode arrays (CFMAs) are described in detail elswhere^41–43,53^. Briefly, a printed circuit board (PCB) was custom manufactured (MicroConnex, Snoqualmie, WA, USA). A connector (A79024-001, Omnetics Connector Corp., Minneapolis, MN, USA) was soldered on one end of the PCB and covered with epoxy. On the other end, 16 bare carbon fibers (T-650/35 3 K, Cytec Industries, Woodland Park, NJ, USA) with a length of 2-3 mm were attached to the PCB in a 2-row (2×8) configuration. The pitch was 132 μm within a row and the separation between the two rows was 50 μm. The array was coated with approximately 800 nm of parylene-c (PDS 2035, Specialty Coating Systems Inc., Indianapolis, IN, USA) for insulation. The insulated carbon fibers had a diameter of 8-9 μm and were cut down to 150-250 μm in length. The base of the carbon fibers were submerged in water and the tips were sharpened with a blowtorch^30^ (MT-51, Master Appliance Corp., Racine, WI, USA) after aligning the carbon fibers with their reflection on the underside of the water surface. The exposed carbon on the sharpened tips (135-160 μm) were electrodeposited with poly(3,4-ethylene-dioxythiophene):sodium p-toluenesulfonate (PEDOT:pTS) by applying 600 pA/fiber for 600 sec. Finally, individual ground and reference wires (AGT05100, World Precision Instruments, Sarasota, FL, USA) were soldered to the PCB.

### Design of Nerve-Holder

To facilitate the insertion of a CFMA in a vagus nerve, we designed a nerve-holder to secure and elevate the vagus nerve away from fluid and breathing motions of the cervical cavity, and allow accurate positioning of a small camera to visualize the CFMA-nerve interface during insertion.

The nerve-holder had a hollow center to allow insertion of carbon fibers without breakage, and to drain excess fluid around the nerve which may obscure the camera view. To handle the nerve-holder, a circular threaded rod (21YN67, Grainger Inc., Lake Forest, IL, USA) was inserted in the holder and was connected to a soldering arm (900-015, Eclipse, Amelia Court House, VA, USA). The nerve-holder was designed using a computer-aided design (CAD) software (Fusion 360, Autodesk, San Rafael, CA, USA) and 3D-printed with clear resin (Form 2, Formlabs, Somerville, MA, USA). The dimensions of the design are shown in Supplementary Fig. S3.

### Animal Surgery

All experimental procedures were approved by the University of Michigan Institutional Animal Care and Use Committee (IACUC) in accordance with the National Institutes of Health’s guidelines for the care and use of laboratory animals. Non-survival experiments were performed on male (0.48-0.83 kg) and female (0.36-0.42 kg) Sprague-Dawley rats (Charles Rivers Laboratories, Wilmington, MA, USA). The animals were housed in ventilated cages under controlled temperature, humidity, and photoperiod (12-h light/dark cycle), and provided with laboratory chow (5L0D, LabDiet, St. Louis, MO, USA) and tap water ad libitum. The rats were fasted for 3 hours before the procedure. Anesthesia was induced by 5% isoflurane (Fluriso, VetOne, Boise, ID, USA) and maintained at 2-3% isoflurane. Rats were placed on a heating pad (ReptiTherm, Zoo Med Laboratories Inc., San Luis Obispo, CA, USA). A vitals-monitor (SurgiVet, Smiths Medical, Norwell, MA, USA) was used to monitor heart rate with an oxygen saturation (SpO2) sensor. A midline ventral cervical incision was made, and retractors (17009-07, Fine Science Tools Inc., Foster City, CA, USA) were used to maintain the cervical opening. Using a dissection microscope (Lynx EVO, Vision Engineering Inc., New Milford, CT, USA), the left cervical vagus nerve (9-12 mm in length) was isolated from the carotid artery and surrounding tissue using fine forceps (00632-11, Fine Science Tools Inc., Foster City, CA, USA). The vagus nerve was lifted (~2 mm) and placed on the nerve-holder to facilitate CFMA insertion. The heating pad and dissection microscope were disconnected to reduce electrical noise.

### CFMA Insertion

The CFMA was accurately controlled by a micromanipulator (KITE-R, World Precision Instruments, Sarasota, FL, USA) that was secured on an optical breadboard (MB1218, Thorlabs Inc., Newton, NJ, USA) under the animal. The ground wire for the CFMA was inserted subcutaneously in the cervical region and the reference wire was placed in fluid or tissue underneath the nerve-holder. A small pen-shaped camera (MS100, Teslong, Shenzhen, China) was placed in the cervical opening to visualize and align the CFMA fibers for insertion. The nerve was rinsed with saline (0.9% NaCl, Baxter International Inc., Deerfield, IL, USA) and the CFMA was inserted in the vagus nerve.

The CFMA was connected to a neural interface processor (Grapevine, Ripple LLC, Salt Lake City, UT, USA) that recorded signals at a sampling rate of 30 kHz on a linked desktop computer. Impedances were measured with the neural interface processor at 1 kHz in saline before the procedure, in the nerve immediately after insertion, and in the nerve at the end of the experiment.

### Experimental Protocol

After completion of surgery and insertion of the CFMA, a baseline recording period of at least 5 minutes was obtained. A dose of glucose (n=6; 1 g, Dextrose 50%, Hospira, Lake Forest, IL, USA), insulin (n=6; 20 U, Vetsulin, Merck Animal Health, Madison, NJ, USA), 2-deoxy-D-glucose (n=6; 60 mg, D8375-1G, Sigma-Aldrich, St. Louis, MO, USA), or saline (n=4; 1.0 mL, 0.9% NaCl, Baxter International Inc., Deerfield, IL, USA) was injected intraperitoneally (IP). Recordings from the CFMA were continued for 60 minutes after the injection. Measurements of blood glucose concentration with a glucometer (AlphaTRAK 2, Abbott, Abbott Park, IL, USA), heart rate with the SpO2 sensor, and breathing rate with a timer were obtained every 5 minutes. The glucometer was unable to measure blood glucose concentrations above 750 mg/dL in one experiment due to the limitations of the glucometer. In experiments with observed breathing-related neural signals (n=3), a recording period of 360 seconds was obtained starting at 2% isoflurane for 60 seconds, followed by 300 seconds at 5% isoflurane. Breathing rate measurements were taken at 30, 150, and 300 seconds from the start of the recording. At the end of the experiment, animals were euthanized with an overdose of sodium pentobarbital (400 mg/kg IP, Euthanasia Solution, VetOne, Boise, ID, USA).

### Analysis of Neural Recordings

The recorded signals were sorted into clusters using Wave_clus^54^, which is a spike-sorting MATLAB-based algorithm that uses wavelet decomposition to extract waveform features and superparamagnetic clustering to cluster the spikes. The signals were filtered with a band-pass filter at 300-10,000 Hz. The minimum spike detection threshold was set between 3.3 and 10.1σ and the maximum threshold was set at 50σ [σ = median(|filtered signal| / 0.6745)]^54^. The sorted clusters were exported to MATLAB (R2014b, MathWorks, Natick, MA, USA) for analysis. Firing rates were calculated with a bin duration of 1 sec. To calculate signal-to-noise ratio (SNR), the mean peak-to-peak amplitude (Vpp) of a sorted cluster was determined and noise intervals with a total duration of at least 7 sec were specified at periods with no occurring spikes or artifacts [SNR = Vpp / (2 x standard deviation of noise)]^40,42^. Cross-correlation was performed between the sorted clusters across all the recording carbon fibers of a CFMA to inspect the latency of spikes along the CFMA. Latencies with high occurrences (count >> mean occurrence) were identified, and the signal traces on adjacent recording carbon fibers were manually reviewed to confirm instances of signal propagation before determining the conduction velocity and signal propagation direction for these high-occurring latencies. The bin-size for the latency counts was set at 0.2 msec, except for one experiment that had multiple high counts at zero latency with this 0.2 msec bin-size resolution. For this experiment only, the bin-size was set at 0.01 msec to provide latency counts with higher resolution. The correlation coefficient (ρ) was calculated for all sorted clusters between the firing rate of each cluster and the measured physiological parameters (breathing rate, heart rate and blood glucose concentration) for that experiment. Since the physiological measurements were much less frequent (every 5 minutes) than cluster firing rates (every second), the average cluster firing rate for 1 minute, centered at the time of each physiological measurement, was determined and used for the correlation coefficient computations. For clusters that responded to a glucose modulation dose (administered at 450 sec), the mean firing rate was calculated before the dose (0-400 sec), and after the dose (500-3000 sec) for comparison. When appropriate, data are presented as mean ± standard deviation (SD).

## Data Availability

All raw recordings, sorted neural clusters, and analysis codes will be available on the Blackfynn data repository platform at DOI: https://doi.org/10.26275/j5wc-rwcr once it has completed NIH SPARC data curation.

## Acknowledgements

We thank Dr. Randy Seeley and Dr. Malcolm Low for their advice on designing the experiments, Dr. Zach Sperry, Aileen Ouyang, Eric Kennedy and Dr. Lauren Zimmerman for their assistance in surgical preparation, Joey Letner for his guidance on data analysis, and Dr. Steve Kemp, Dr. Dan Ursu, Jana Moon, Charles Hwang and Sasha Meshinchi for preparing and imaging the immunohistochemistry nerve sample. This research was supported by the National Institutes of Health (NIH) Stimulating Peripheral Activity to Relieve Conditions (SPARC) Program (Award 1OT2OD024907) and the National Science Foundation (Award 1707316).

## Author Contributions

Planned study – AAJ, JPS, CAC, TMB. Fabricated arrays – EJW, PRP, JMR, CAC. Performed surgeries and collected data – AAJ, DCR, ECB, TMB. Analyzed data – AAJ, DCR, ECB, TMB. Drafted manuscript – AAJ, TMB. Reviewed manuscript and approved final version – AAJ, DCR, EJW, PRP, JMR, ECB, JPS, CAC, TMB.

