## Supplementary Figures S1-S3 for "Multi-channel intraneural vagus nerve recordings with a novel high-density carbon fiber microelectrode array"

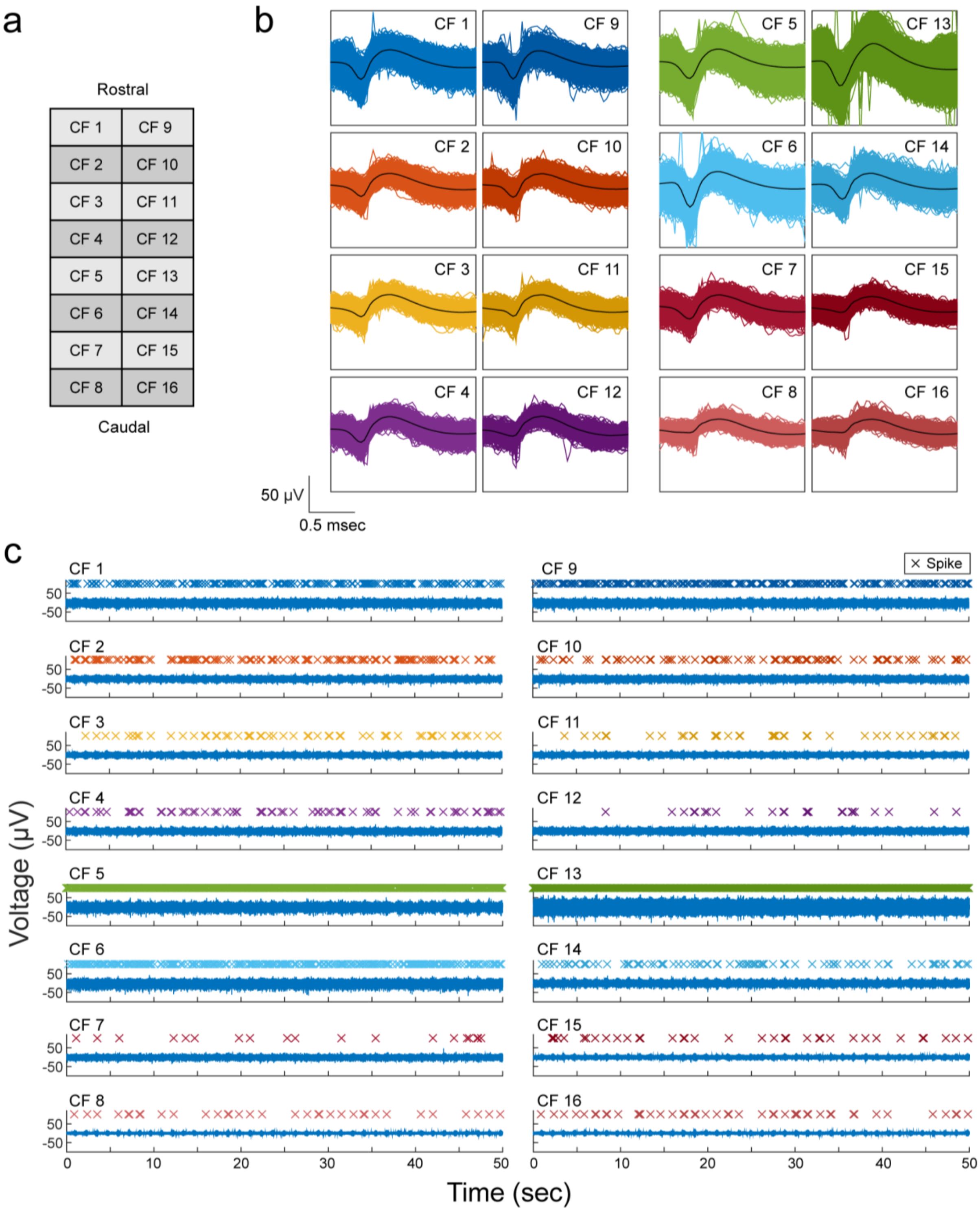


**Supplementary Figure S1. Carbon Fiber Microelectrode Array (CFMA) with neural activity on all recording carbon fibers (CFs). (a)** The CF layout in a CFMA. **(b)** Sorted neural cluster on each CF. **(c)** Segment of filtered signal on each CF. The firing of spikes are unique across CFs.

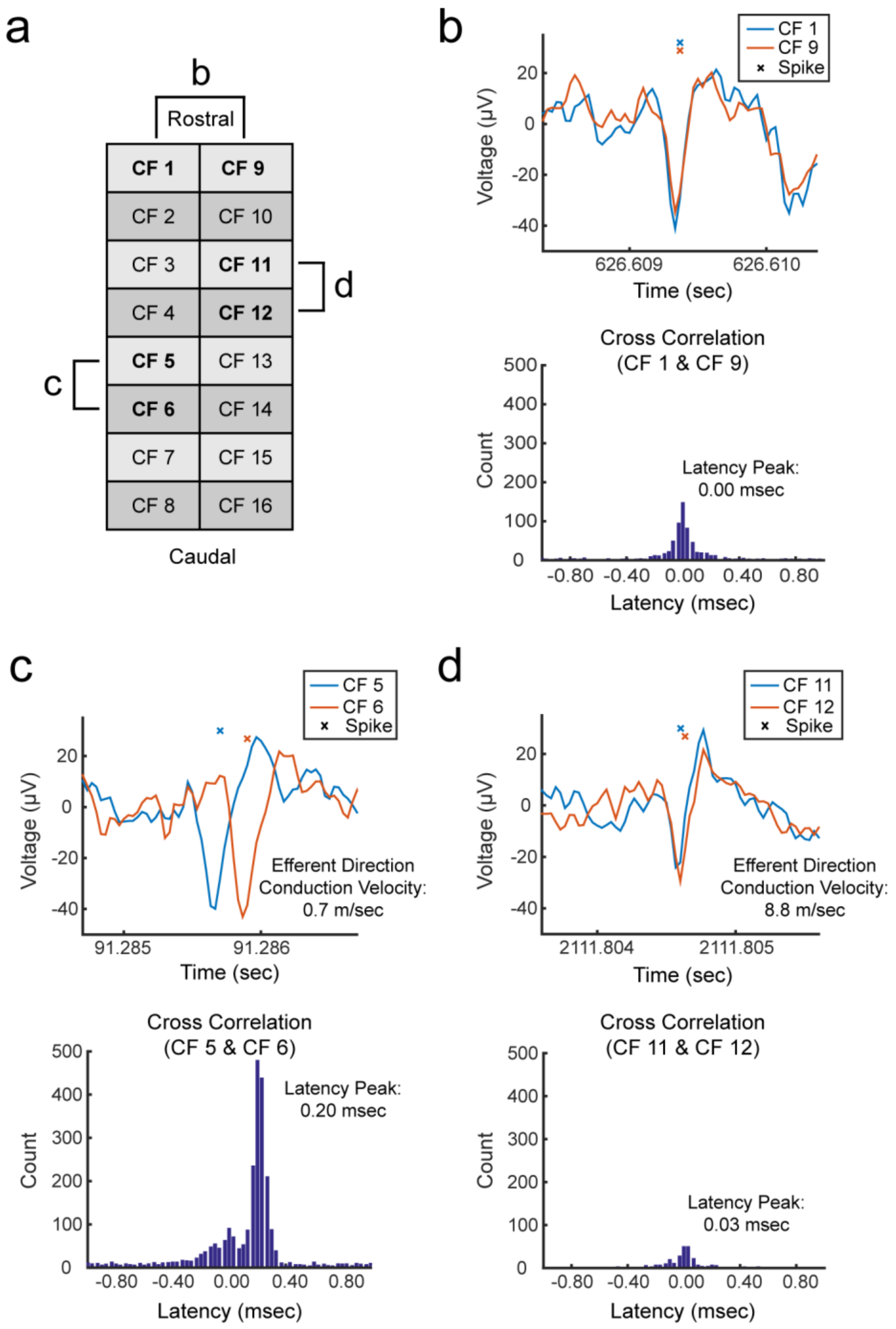


**Supplementary Figure S2. Neural recording on opposite rows of a CFMA. (a)** The 2-row carbon fiber (CF) configuration in a CFMA. **(b)** Two CFs on opposite rows showing coinciding spikes with different amplitudes, suggesting that these spikes are generated from a neuron located between these opposite carbon fibers. **(c)** Adjacent carbon fibers showing propagating spikes in the efferent direction at a conduction velocity of 0.7 m/sec along one row of a CFMA. (d) Propagating spikes on the other row with a conduction velocity of 8.8 m/sec in the efferent direction. The propagating spikes in c and d are independent of each other.

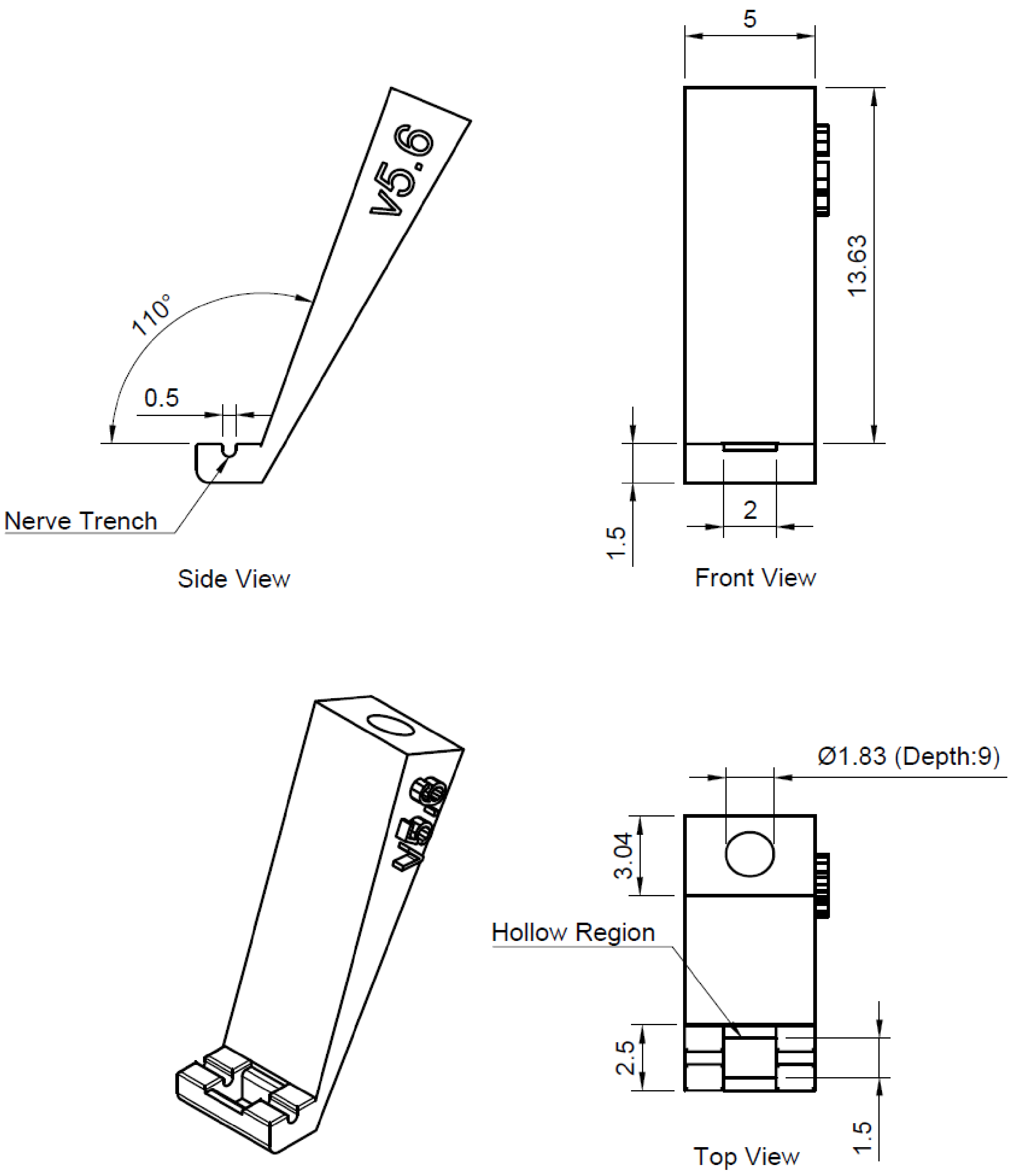


**Supplementary Figure S3. Nerve-holder design for inserting CFMA in a rat cervical vagus nerve.** Dimensions are in millimeters (mm).
